# Polymerization of misfolded protein lowers CX3CR1 expression in human PBMCs

**DOI:** 10.1101/2020.11.24.395640

**Authors:** Srinu Tumpara, Matthias Ballmaier, Sabine Wrenger, Mandy König, Matthias Lehmann, Ralf Lichtinghagen, Beatriz Martinez-Delgado, David DeLuca, Nils Jedicke, Tobias Welte, Malin Fromme, Pavel Strnad, Jan Stolk, Sabina Janciauskiene

## Abstract

The CX3CR1 (chemokine (C-X3-C motif) receptor 1) expression levels on immune cells have significant importance in maintaining tissue homeostasis under physiological and pathological conditions. The factors implicated in the regulation of CX3CR1 and its specific ligand CX3CL1 (fractalkine) expression remain largely unknown. Recent studies provide evidence that host‘s misfolded proteins occurring in the forms of polymers or amyloid fibrils can regulate CX3CR1 expression. Herein, we present a novel example that polymers of human ZZ alpha1-antitrypsin (Z-AAT) protein, resulting from its conformational misfolding due to the Z (Glu342Lys) mutation in *SERPINA1* gene, strongly lower CX3CR1 expression in human PBMCs. We also show that extracellular polymers of Z-AAT are internalized by PBMCs, which parallels with increased intracellular levels of CX3CR1 protein. Our findings support the role of extracellular misfolded proteins in CX3CR1 regulation and encourage conducting further studies on this issue.

## Introduction

The interaction between the chemokine receptors and chemokines, but also other proteins, peptides, lipids, and microbial products, plays a critical role in the recruitment of inflammatory cells into injured/diseased tissues^1^. Many human diseases involve altered surface expression of chemokine receptors, which can lead to a defective cell migration and inappropriate immune response. Most of the human PBMCs express CX3CR1^2^, also known as the G-protein coupled receptor 13 (GPR13) or fractalkine receptor, a mediator of leukocyte migration and adhesion. In the central nervous system, CX3CR1 is largely expressed by microglial cells (brain macrophages)^3^, which play a critical role in the progression of neurodegenerative diseases like Alzheimer’s disease. The major role of CX3CR1-expressing cells is to recognize and enter tissue following CX3CL1 (fractalkine or also called neurotactin) gradient, and to crawl or “patrol” in the lumen of blood vessels^4^. Since CX3CR1/CX3CL1 axis is also involved in the synthesis of anti-inflammatory cytokines and has a significant role in cytoskeletal rearrangement, migration, apoptosis and proliferation, its dysregulation is associated with the development of cardiovascular diseases, kidney ischemia–reperfusion injury, cancer, chronic obstructive pulmonary disease, neurodegenerative disorders and others^5, 6, 7^. Some studies indicate that CX3CR1 deficiency contributes to the severity of infectious diseases^8^, and promotes lung pathology in respiratory syncytial virus-infected mice^9^. It is shown that animals with deletion of CX3CR1 have impairments in phagocytosis^10^, which is vital to prevent unwanted inflammation. It is clear that CX3CR1 expressing cells have tissue-specific roles in different pathophysiological conditions. Nevertheless, a comprehensive knowledge on the regulation of CX3CR1 expression is still missing.

One of the important aspects that needs to be addressed in the chemokine-receptor regulation is the ability of ligands to induce internalization of the receptors and to internalize themselves during this process. Current findings suggest that divergent proteins with a common propensity to form extracellular polymers, independently of amino acid sequence, size, structure, expression level, or function might interact with chemokine receptors and affect their surface expression levels. For example, it has been reported that Alzheimer‘s peptide, Aβ, interacts with CX3CR1, and significantly reduces its expression in cultured microglial cells and Alzheimer‘s brain^12^. Similarly, highly accumulated extracellular Tau in Alzheimer‘s disease, seems to bind to CX3CR1 promoting its internalization and reducing expression in microglial cells^13^. In concordance, we provide novel evidence that polymers of human Z alpha1-antitrypsin (Z-AAT), resulting from its conformational misfolding due to the Z (Glu342Lys) mutation in *SERPINA1* gene, strongly lower CX3CR1 expression in human PBMCs. We also show that extracellular oligomers of Z-AAT are internalized by PBMCs, which parallel with increased intracellular CX3CR1 protein levels.

## Results and Discussion

Inherited α1-antitrypsin deficiency (AATD) is a rare genetic condition caused by mutations in the *SERPINA1* gene. Homozygous Z AATD is the most clinically relevant genotype among Caucasians (prevalence is about 1:2000-1:5000) that is characterized by low plasma levels of AAT protein (10-15% compared to the wild type, MM AAT, 1.3-2 g/L) and the presence of intracellular and circulating Z-AAT polymers^14^.

The intracellular polymers of Z-AAT are known to be harmful for AAT-producing cells whereas in the circulation these polymers are not able to execute the tasks of native AAT-a major inhibitor of neutrophil serine proteases having broad spectrum anti-inflammatory activities. Based on the fact that circulating Z-AAT polymers are dysfunctional and contribute to the risk of developing lung and/or liver pathology^15, 16^, we wanted to investigate CX3CR1 expression in PBMCs of ZZ AATD individuals. For this, in collaboration with German Alpha1 Patient Association and Aachen University we prepared RNA from PBMCs isolated from 41 clinically stable ZZ AATD volunteers independently of their clinical diagnosis or treatment with intravenous AAT, a specific augmentation therapy^17^. For comparison, we used total PBMCs from volunteers having normal plasma AAT levels. Additionally, a limited amount of RNA sample was available from PBMCs isolated from a cohort of 12 ZZ AATD emphysema patients at Leiden University Medical Center (**Figure 1–figure supplement 1**). To our surprise, the *CX3CR1* mRNA expression turned to be much lower in ZZ AATD PBMCs than in PBMCs from non-AATD controls [median (range): 4.1 (2.7-5.5) vs 18.5 (13-26.6), p < 0.001] (**Figure 1A**) independent of individual’s age, clinical diagnosis (healthy, lung or liver disease) or augmentation therapy. A previous study has shown that CX3CR1^-/-^ mice have significantly higher plasma levels of CX3CL1 than wild-type mice^18^ and that CX3CL1 reveals its biological activity exclusively via an interaction with CX3CR1^1^. Therefore, we thought that the lower expression of *CX3CR1* by ZZ AATD PBMCs might be associated with attenuated levels of soluble CX3CL1. However, plasma levels of CX3CL1 were low and did not differ between ZZ AATD and non-AATD individuals (**Figure 1B**), and did not correlate with CX3CR1 expression in PBMCs. The expression and release of CX3CL1 is generally low in the absence of inflammatory insults^18^ showing that at the time point of blood donation all volunteers were under stable clinical condition.

**Figure 1.**
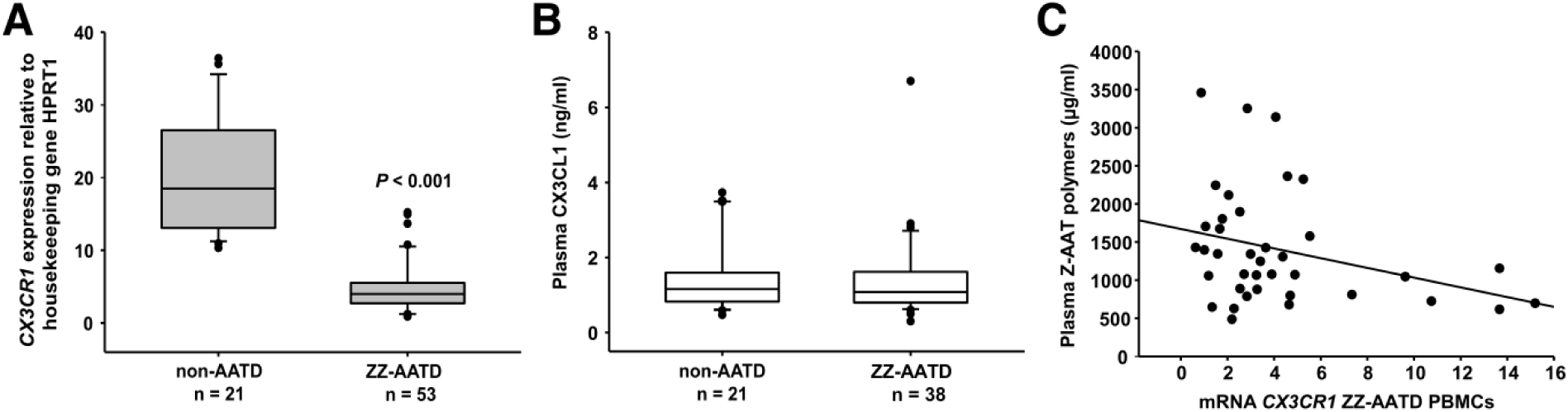
**A** *CX3CR1* gene expression in PBMCs isolated from AATD subjects and non-AATD controls. *CX3CR1* gene expression relative to *HPRT1* housekeeping gene was determined by real-time PCR using Taqman gene expression assays. Measurements were carried out in duplicates. Data are presented as median (IQR) in boxplots, lines represent medians. Outliers are defined as data points located outside the whiskers. p-value was calculated by the difference between the Mann-Whitney U test. **B** Plasma levels of CX3CL1 in AATD (plasma available for n = 38 AATD) and non-AATD individuals measured by ELISA. Measurements were carried out in triplicates. Data are presented as median (IQR) in boxplots with whiskers. Outliers are defined as data points located outside the whiskers. **C** Negative correlation of *CX3CR1* mRNA in PBMCs and plasma Z-AAT polymer levels from ZZ-AATD individuals from graph (**B**). Person’s correlation test, r^2^ = -0.313, p = 0.055, n = 38.

CX3CR1 is preferentially expressed on CD14^+^ monocytes and NK cells^20^, and previous reports indicated that exogenous IL-15 is a negative regulator of CX3CR1 expression in human CD56^+^ NK cells^21, 22^. According to our data, plasma levels of IL-15 were lower in ZZ AATD than in non-AATD (pg/ml, median (range): 6.6 (5.9-6.9), n = 23 vs 7.63 (6.63-8.1), n = 21, p = 0.001). Thus, we excluded a possible link between exogenous IL-15 and expression of *CX3CR1* in PBMCs.

Taken into account that ZZ AATD individuals, differently from non-AATD, have about 90% lower blood concentration of Z-AAT protein, which may affect cellular microenvironment^23^ and the expression of CX3CR1, we thought that there might be a relationship between gene expression of *CX3CR1* and Z-AAT plasma levels. However, we could not find any correlation between *CX3CR1* mRNA in PBMCs and plasma levels of Z-AAT measured by nephelometry (data not shown). Therefore, we next decided to measure plasma Z-AAT polymers because Z mutation destabilises AAT protein and causes the formation of polymers that are present in the circulation of all carriers of the Z allele^14, 24^. As anticipated, only minor amounts of polymers were detected in plasma of non-AATD individuals while plasma of ZZ AATD contained high amounts of polymers [µg/ml, mean (SD): 4.1 (6), n = 18 vs 1399.8 (750), n = 20, respectively]. Since most of the ZZ AATD individuals received intravenous augmentation therapy with plasma purified wild-type AAT protein, we segregated ZZ AATD individuals into subgroups who receive or not receive therapy but found no statistical difference in Z-AAT polymer levels [µg/ml, median (range): non-augmented 1506.6 (854-178.1), n = 17 vs augmented 1348.5 (779.5-152.9), n = 23, respectively]. Hence, we confirmed that ZZ AATD people have high levels of plasma Z-AAT polymers, independent on the therapy with AAT preparations. It is important to point out that a previous study using sandwich ELISA based on 2C1 antibody found that circulating Z-AAT polymers range between 8.2-230.2 μg/ml in ZZ AATD^14^ whereas by using our single monoclonal antibody (LG96)-based ELISA we detect much higher circulating levels of Z-AAT polymers. These discrepancies can be due to the differences between antibody specificities. For example, 2C1 showed high affinity for polymers formed by heating M or Z AAT at 60°C ^25^ while LG96 antibody recognizes naturally occurring/native Z-AAT polymers without requiring sample heating. Indeed, to provide further explanation why in some individuals serum levels of Z-AAT polymers are higher than Z-AAT monomers (measured by nephelometry) is of great importance and needs to be investigated in more detail in future studies.

Most interestingly, however, we found a trend towards an inverse relationship between *CX3CR1* mRNA and plasma Z-AAT polymers (r^2^ = -0.31, n = 38, p = 0.055) (**Figure 1C**). This latter prompted us to investigate whether Z-AAT polymers isolated from ZZ AATD individuals affect CX3CR1 expression when added to healthy donor PBMCs, *in vitro*. We incubated PBMCs obtained from healthy donors either alone or with addition of Z-AAT, or M-AAT for 18 h. LPS (1 µg/mL) was included as a known reducer of CX3CR1 expression^26,27^. Indeed, polymeric Z-AAT in a concentration-dependent manner lowered *CX3CR1* mRNA expression (**Figure 2–figure supplement 2**) whereas repeated experiments using Z-AAT at a constant concentration of 0.5 mg/mL reduced *CX3CR1* expression more than twice as compared to non-treated controls (**Figure 2A**). In accordance with the gene expression results, LPS and polymer containing Z-AAT preparation significantly decreased surface expression of CX3CR1, specifically in CD14^+^ monocytes and NK cells (**Figure 3A**,**B**). Remarkably, western blot analysis revealed high levels of CX3CR1 protein in the presence of Z-AAT and LPS (used as a positive control) in total cell lysates (**Figure 2B**,**C**). In addition, CX3CR1 protein presented in detergent resistant lipid rafts of PBMCs treated with Z-AAT (**Figure 2–figure supplement 3A**). Total cell lysates and lipid raft fractions from Z-AAT treated PBMCs contain high amounts of polymers compared to cells treated with M-AAT preparations (**Figure 2C, figure 2-figure supplement 3B**). Under the same experimental conditions, M-AAT had no effect on CX3CR1 mRNA [expression relative to housekeeping gene HPRT1, mean (SD): 24.9 (2.9) controls, n=5 vs 23.7 (1.3), n=5, NS], CX3CR1 surface expression and cellular protein levels (**Figure 2B**).

**Figure 2.**
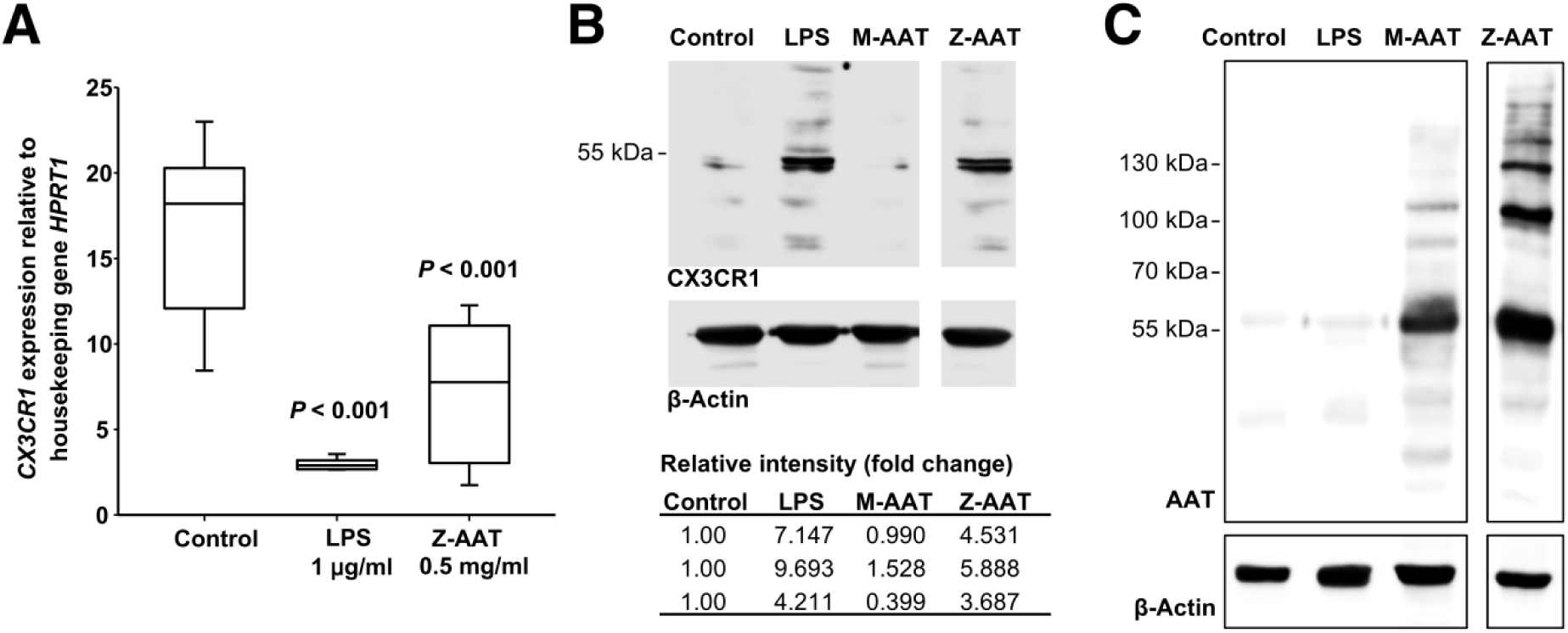
Inverse effects of Z-AAT on *CX3CR1* gene expression and protein expression. PBMCs were incubated for 18 h with plasma-derived Z-AAT, LPS or M-AAT in the concentrations as indicated, or with RPMI medium alone (control). **A** *CX3CR1* gene expression relative to *HPRT1* housekeeping gene was determined by real-time PCR using Taqman gene expression assays. The data from n = 6 independent experiments are presented as median (IQR) in box and whisker plot format; lines represent medians in each box. Measurements were carried out in duplicates. p was calculated by nonparametric Kruskal-Wallis test). **B** Representative uncut Western blot (n = 3 independent experiments) of CX3CR1 in RIPA lysates prepared from PBMCs incubated for 18 h alone or with LPS (1 µg/ml), M-AAT (1 mg/ml), or Z-AAT (0.5 mg/ml). For analysis of CX3CR1, equal amounts of protein were separated by SDS-PAGE under reducing conditions. Relative intensities were calculated for each band using the ratio relative to β-actin, as a loading control, and then normalized by the experimental control. (Table, n = 3 independent experiments). **C** For analysis of cellular AAT the same lysates were separated under non-reducing conditions. Western blots were probed with polyclonal rabbit anti human AAT recognizing monomeric, polymeric, or truncated forms of AAT. One representative blot from n = 3 independent experiments is shown. β-actin was used for a loading control.

**Figure 3.**
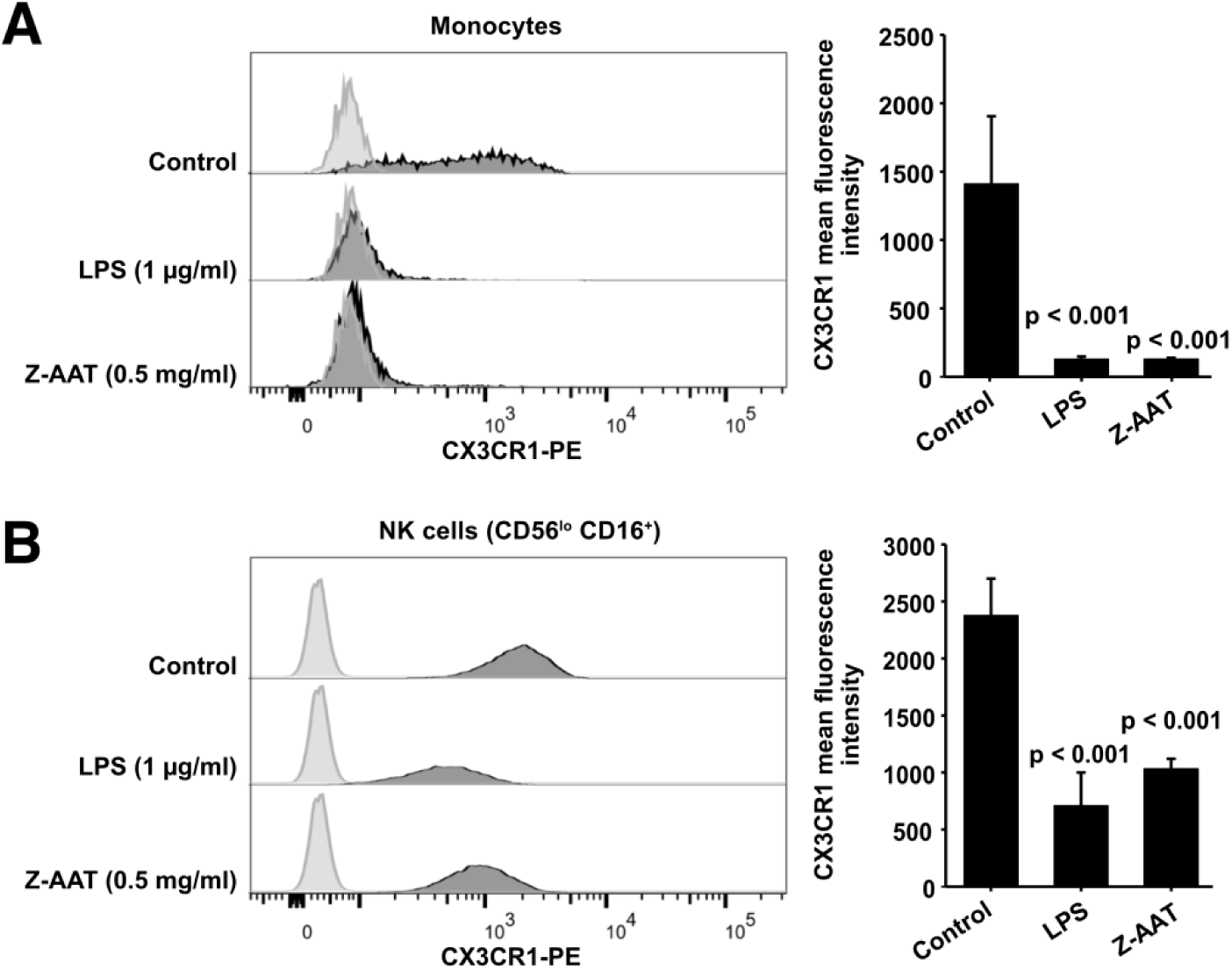
Flow cytometric analysis of CX3CR1 surface expression in PBMCs after incubation with RPMI alone (control), Z-AAT or LPS for 18 h. CX3CR1 expressing cells were found in the monocyte gate (**A**) and in the NK cell gate (CD56^lo^ CD16^+^) (**B**). Histograms show representative results and bars represent mean (SD) of n = 4 independent biological repeats each measured one time. After incubation with Z-AAT or LPS monocytes and NK cells show significantly reduced CX3CR1 surface expression in comparison to untreated control cells. p-values were calculated by one-way ANOVA.

Altogether, our data show that polymeric Z-AAT, similar like LPS, decreases CX3CR1 mRNA and cell surface expression but instead increases an intracellular pool of CX3CR1 which parallels with increased intracellular polymeric Z-AAT. In general, along with transcriptional regulation, translocation into the cell membrane is important for the regulation of chemokine receptors^28^. To achieve a definitive answer how Z-AAT or other types of protein polymers regulate CX3CR1 expression detailed mechanistic studies are required.

After *in vitro* challenge with LPS for longer periods (like for 18 h), human monocytes are known to increase in the mRNA and membrane expression of CD14, a receptor for LPS^29^. The enhancement of CD14 expression after treatment of PBMCs with Z-AAT strikingly resembled that induced by LPS (**Figure 4A**,**B** and **Figure 4–figure supplement 1**). This raised a suspicion that Z-AAT preparations might contain endotoxin. According to the Limulus Amebocyte Lysate Test (LAL), endotoxin levels of Z-AAT preparations were below detection limit (0.01 EU/ml). Moreover, considering the importance of CD14 in regulating pro-inflammatory cytokine production, we measured expression and release of pro-inflammatory cytokines in PBMCs treated with LPS or Z-AAT. LPS significantly induced expression of TNFα, IL-6 and IL-1β while polymer containing Z-AAT preparations did not stimulate the expression of these cytokines (**Figure 4 –figure supplement 2**). In line, LPS but not Z-AAT significantly increased release of cytokines [IL-1β, pg/ml, median (range): LPS 1342.9 (1008-1834) vs Z-AAT 3.2 (2.5-5.9) vs controls 2.5 (2.1-3.7), n = 4 independent experiments; TNFα, ng/ml, mean (SD): LPS 19.5 (2.5) vs Z-AAT vs controls, undetectable, n = 4 and IL-6, ng/ml, median (range): LPS 15903.5 (14626-17262) vs Z-AAT 5.4 (2.9-6.1) vs control (1.7 (1.0-2.3), n = 4]. Therefore, we concluded that the effect of Z-AAT preparations on CD14 is valid and unrelated to a potential LPS-contamination.

**Figure 4.**
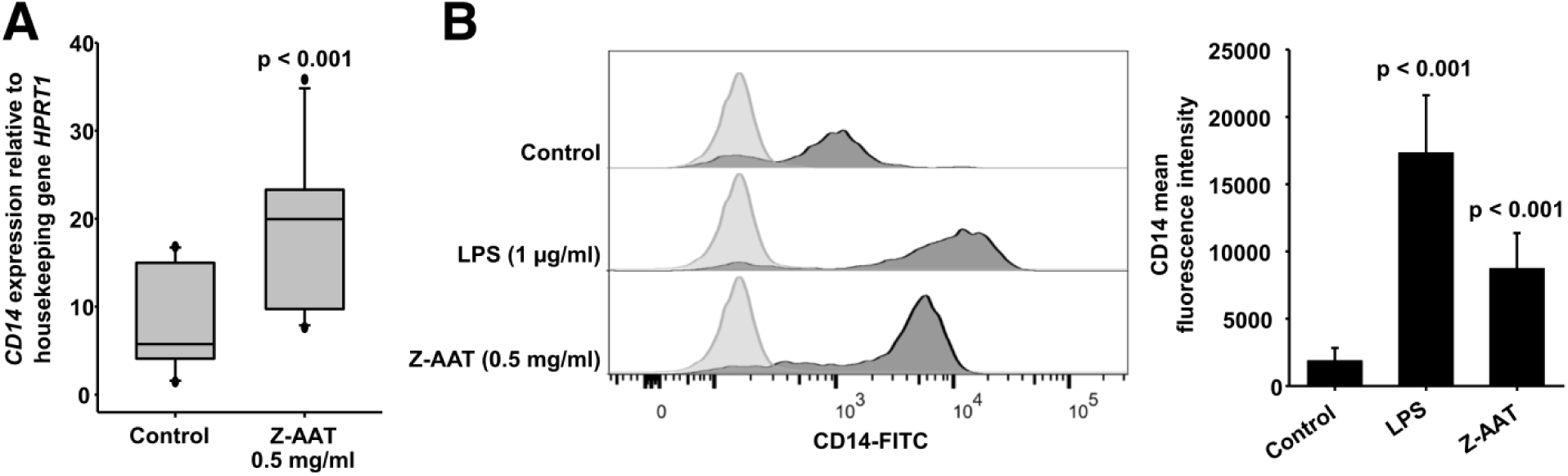
Z-AAT and LPS induce CD14 expression. **A** Z-AAT increases *CD14* gene expression. PBMCs were incubated for 18 h with 0.5 mg/ml Z-AAT or RPMI medium alone (control). *CD14* mRNA expression relative to HPRT1 was determined by real-time PCR using Taqman gene expression assays. Measurements were carried out in duplicates. Data are represented as median (IQR) in boxplots, lines represent medians of n = 14 independent biological repeats. Outliers are defined as data points located outside the whiskers. p-values were calculated using nonparametric Kruskal-Wallis test. **B** Z-AAT increases monocyte CD14 surface expression. PBMCs were cultured with RPMI (control), Z-AAT or LPS for 18 h. CD14 mean fluorescence intensities of monocytic cells were determined by flow cytometry. Histograms show representative results and bars represent mean (SD) of n = 4 independent biological repeats each measured one time. p-values were calculated from ANOVA.

As a side note, it has been reported that CD14^++^ monocytes have the lowest expression of CX3CR1^30^. Low and high surface CX3CR1 levels are suggested to delineate two functional subsets of murine blood monocytes: “inflammatory” and “resident monocytes,”^31^. This dichotomy appears conserved in humans as CD14^+^ CD16^−^, and CD14^low^ CD^16+^ monocytes resemble “inflammatory” and “resident” monocytes. We previously demonstrated that peripheral blood monocytes from clinically healthy young adults (30-year old) with ZZ AATD have significantly higher mRNA and surface expression of CD14 as compared to age matched MM subjects^32^. We thought that the higher CD14 expression reflects early pathological processes whereas according to our current findings this phenomenon seems to relate with the circulating Z-AAT polymers.

In summary, we report a novel observation that polymeric Z-AAT lowers the expression of CX3CR1 on PBMCs. We predict that the relationship between the circulating amounts of Z-AAT polymers and CX3CR1 expression levels in immune cells may account for the individual susceptibility or course of certain diseases. During steady state, monocytes expressing CX3CR1 patrol healthy tissues through crawling on the resting endothelium but these monocytes are required for a rapid tissue invasion at the site of infection or inflammation^33, 34^. Therefore, diminished numbers of CX3CR1-positive cells may favour the development of tissue injury and disease. To date, many functional aspects of the CX3CR1-CX3CL1 axis have been suggested, including the adhesion of immune cells to vascular endothelial cells, chemotaxis, the crawling of the monocytes that patrol on vascular endothelial cells, the retention of monocytes of the inflamed endothelium to recruit inflammatory cells, and the survival of the macrophage. Considering the above, these different aspects of interactions between PBMCs and Z-AAT or other polymers occurring due to genetic or post-translational protein modifications require further investigations in dedicated clinical and experimental studies. We strongly believe that our new observation in general will encourage more detailed analyses of circulating polymers of misfolded proteins and their role in CX3CR1-CX3CL1 regulation during health and diseases.

## Material and methods

### Study approval

The study cohort consists of 41 clinically stable ZZ AATD volunteers collected in collaboration with German Alpha1 Patient Association and Aachen University independently on their clinical diagnosis or treatment with intravenous AAT and 21 non-AATD healthy controls. For Z-AAT polymer determination, we added 12 ZZ AATD emphysema patients recruited at Leiden University Medical Center. The institutional review board of Aachen University (EK 173/15) provided ethical approval for individuals recruited in Germany. Leiden University Medical Center provided ethical approval (project P00.083 and P01.101) for the second study group. For all individuals detailed medical records data were anonymized. All participants issued a written informed consent according to the ethical guidelines of the Helsinki Declaration (Hong Kong Amendment) as well as Good Clinical Practice (European guidelines).

### Isolation of PBMCs

Total PBMCs were isolated from freshly obtained peripheral blood (within 6 hours) using Lymphosep (C-C-Pro, Oberdorla, Germany) discontinuous gradient centrifugation according to the manufacturer’s instructions as described previously^35^. Thereafter, cells were lysed with RLT buffer for RNA analysis or suspended in RPMI-1640 medium (Gibco, Thermofisher Scientific, Waltham, MA, USA) and plated into non-adherent 12-well plates (Greiner Bio-one, Kremsmünster, Austria) for the further analyses.

### RT-PCR analysis

Isolation of total RNA, synthesis of cDNA and mRNA analysis using Taqman gene expression assays (Thermofisher Scientific, Waltham, MA, USA, **Table 1**) were performed as described previously^35^. RT PCR was carried out in duplicates. RNA quality was checked on agarose gels.

**Table 1.** Primers for gene expression analysis.

| Primer | Assay ID |
| --- | --- |
| **CX3CR1** | Hs00365842_m1 |
| **TNFA** | Hs01113624_m1 |
| **IL6** | Hs00985639_m1 |
| **IL1B** | Hs01555410_s1 |
| **HPRT1** | Hs02800695_m1 |
| **CD14** | Hs00169122_g1 |

### AAT polymer ELISA

The AAT polymer ELISA using the monoclonal antibody LG96 (deposited under access No DSM ACC3092 at German Collection of Microorganisms and Cell Cultures) was developed by Candor Biosciences. Normal M-AAT was used for a negative control. Recovery ratio, signal-to noise ratio, calibration curve, sample stability under different storage conditions were tested and all tests passed. A cross reactivity with M-AAT was not reported in any of the tests. Nunc MaxiSorp flat-bottom 96-well plates (Thermofisher, Waltham, MA, USA) were coated overnight at 2-8 °C with monoclonal antibody LG96, at 1 µg/ml in coating buffer pH 7.4 (Candor Biosciences, Wangen, Germany). After a 2 h blocking step, the plasma samples were applied in the previously determined dilutions made in LowCross-Buffer (Candor Biosciences), which also served as a blank. Incubation was performed for 2 h at RT. For detection, the captured antigen was incubated with antibody (LG96)-horseradish peroxidase (HRP) conjugate (1:2000) for 2 h. The conjugate was prepared in advance with the HRP Conjugation Kit Lightning-Link (Abcam, Cambridge, UK) according to the manufacturer’s instructions. For signal development SeramunBlau fast2 microwell peroxidase substrate (Seramun, Heidesee, Germany) was used. The incubation was performed at room temperature for 12 minutes in the dark and the reaction was stopped with 2 M H_2_SO_4_. Plates were analyzed at 450 nm by microplate reader (Dynex Chantilly, VA, USA) equipped with Dynex Revelation 4.21 software. Measurements were carried out in triplicates.

### Preparation of AAT proteins

Plasma M- and Z-AAT was isolated by affinity chromatography using the AAT specific Alpha-1 Antitrypsin Select matrix (GE Healthcare Life Sciences, Cytiva, Sheffield, UK) according to the manufacturer’s recommendations. For Z-AAT preparation we pooled plasma from volunteers not receiving AAT augmentation therapy. To change the buffer in the M- and Z-AAT protein pools to Hank’s balanced salt solution (HBSS, Merck, Darmstadt, Germany) we used Vivaspin centrifugal concentrators with 10,000 MWCO (Vivaproducts, Littleton, MA, USA). Plasma purified human AAT (99% purity, Respreeza, Zemaira, CSL Behring, Marburg, Germany) was changed to HBSS by the same method. Protein concentrations were determined using Pierce BCA Protein Assay Kit (Thermofisher, Waltham, MA, USA). The quality of the M- and Z-AAT preparations was confirmed on Coomassie gels (10 % SDS PAGE, **Figure 2 –figure supplement 1**) and by analyzing endotoxin levels with Pierce Chromogenic Endotoxin Quant Kit according to the manufacturer’s guidelines (Thermofisher, Waltham, MA, USA) using TECAN Infinite M200 PRO (Männedorf, Switzerland). In both, M and Z-AAT preparations, endotoxin levels were below the detection limit (Assay sensitivity: 0.01-0.1 EU/ml).

### In vitro experiments with PBMCs from healthy donors

PBMCs (5 × 10^6^ cells/ml) were incubated for 18 h at 37 °C, 5% CO_2_ either alone, or with Z- or M-AAT proteins, or lipopolysaccharide (LPS, 1 µg/ml, *Escherichia coli O55:B5*, Sigma-Aldrich, Merck, St. Louis, Missouri, USA). Afterwards, cells were used for RNA isolation, flow cytometry or Western blot analysis. For Western blot, PBMCs were lysed in RIPAm buffer (Sigma-Aldrich), supplemented with protease inhibitor cocktail (Sigma-Aldrich). For some Western blot experiments, we extracted detergent resistant lipid raft associated proteins from insoluble cell fractions using UltraRIPA kit according to the supplier’s instructions (BioDynamics Laboratory, Tokio, Japan).

### Western blot

Equal amounts of lysed proteins were separated by 7.5% or 10% SDS-polyacrylamide gels (under reducing conditions for CX3CR1 and non-reducing for total AAT or AAT polymer analysis) prior to transfer onto polyvinylidene difluoride (PVDF) membranes (Meck-Millipore, Burlington, MA, USA). Membranes were blocked for 1 h with 5% low fat milk (Carl Roth, Karlsruhe, Germany) followed by overnight incubation at 4 °C with specific primary antibodies: polyclonal rabbit anti-human AAT (1:800) (DAKO A/S, Glostrup, Denmark), mouse monoclonal anti-AAT polymer antibody (clone 2C1, 1:500, Hycult Biotech, Uden, The Netherlands), rabbit polyclonal anti-CX3CR1 (1:500, Abcam, Cambridge, UK), or HRP-conjugated monoclonal anti β-actin antibody (1:20,000, Sigma-Aldrich, Merck, St. Louis, Missouri, USA) for a loading control. The immune complexes were visualized with anti-rabbit or anti-mouse HPR-conjugated secondary antibodies (DAKO A/S) and enhanced by Clarity Western ECL Substrate (BioRad, Hercules, CA, USA). Images were acquired by using Chemidoc Touch imaging system (BioRad) under optimal exposure conditions and processed using Image Lab version 5.2.1. software (Bio-Rad). For quantification, the signal intensity of the CX3CR1 protein band in each lane was divided by the corresponding β-actin band intensity (normalization factor or loading control). Afterwards, the normalized signal of each lane was divided by the normalized target signal observed in the control sample to get the abundance of the CX3CR1 protein as a fold change relative to the control.

### ELISA

Plasma samples from 22 ZZ-AATD and 21 non-AATD controls were analyzed for CX3CL1/ Fractalkine using Duoset kit (R&D systems, Minneapolis, MN, USA, assay sensitivity 0.072 ng/ml, detection range 0.2-10 ng/ml). Cell free culture supernatants were analyzed directly or stored at -80°C. ELISA Duoset kits for TNF-α (assay detection range 15.6-1000 pg/ml), IL-1β/IL-1F2 (assay detection range 3,91-250 pg/ml), and IL-6 (assay detection range 9.38-600 pg/ml) were purchased from R&D Systems (Minneapolis, MN, USA) and were used according to the manufacturer’s instructions. Plates were measured on Infinite M200 microplate reader (Tecan, Männedorf, Switzerland). Measurements were carried out in duplicates.

### Flow cytometry analysis

PBMCs (2 × 10^6^ cells per condition) were incubated with LPS (1 µg/ml), M-AAT (1 mg/ml), or Z-AAT (0.5 mg/ml) for 18 h. Staining was performed with phycoerythrin (PE)-conjugated mouse monoclonal anti-CX3CR1 antibody (clone 2A9-1 Invitrogen, Thermofisher Scientific, Carlsbad, CA, USA), fluorescein (FITC)-conjugated mouse monoclonal anti-CD14 antibody (clone TuK4, Life technologies, Thermofisher Scientific, Carlsbad, CA, USA), allophycocyanin (APC)-conjugated mouse monoclonal anti-CD16 antibody (clone 3G8, Immunotools, Friesoythe, Germany), or BV-480 conjugated anti-CD56 mouse monoclonal antibody (Clone NCAM16.2, BD Biosiences, San Jose, CA, USA) alone or in combinations. Dead cells were excluded by a staining with 7-amino-actinomycin D (7-AAD). Samples were measured on a BD FACSAria Fusion machine and analyzed with FlowJo v10 (Becton, Dickinson and Company, Franklin Lakes, NJ, USA). The gating strategy is shown in **Figure 3 –figure supplement 1**.

### Statistics

Data were analyzed and visualized by using Sigma Plot 14.0. One-tailed Student’s t-test was applied to compare two sample means on one variable. When more than two groups were involved in the comparison, one-way ANOVA was used. Data were presented as mean (SD) but if normality test failed, the nonparametric Kruskal-Wallis one-way analysis or Mann-Whitney Rank Sum Test was performed, and data were presented as median (range). For correlation analysis the Pearson’s linear correlation method was used to measure the correlation for a given pair. A p-value of less than 0.05 was considered significant.

## Acknowledgments

We thank the German society Alpha1 Deutschand e.V. and society members for support.

## Additional information

### Author contributions

Conceptualization, SJ, RS, MB, TW, JS, PS; sample collection: NJ, PS, JS; investigations: ST, BMD, SW, RL, MB; data analysis and statistics: ST, BMD, SJ: manuscript preparation: SJ, SW; review and editing: SJ, ST, BMD, RS, SW, NJ, RL, MB, TW, PS, JS.

### Ethics

The institutional review board of Aachen University (EK 173/15) provided ethical approval for individuals recruited in Germany. Leiden University Medical Center provided ethical approval (project P00.083 and P01.101) for the second study group.

### Additional files Supplementary files

Source data and figure supplements are provided in supplementary file *Supplementary file Tumpara et al*.

## Figures and legends

The online version of this article includes the following source data and figure supplement(s) for figure 1:

**Source data 1**. Source files, containing original data for Figure 1 **A**,**B**, to document CX3CR1 expression (**A**), and plasma levels of CX3CR1 in AATD and non-AATD individuals (**B**).

**Figure supplement 1**. Schematic presentation of the study design.

The online version of this article includes the following source data for figure 2: Source data 1. Source files, containing original data for Figure 2**A**, to document CX3CR1 reduced expression in PBMCs treated with Z-AAT or LPS (**A**).

**Figure supplement 1**. Quality control of isolated M- and Z-AAT protein by SDS-PAGE.

**Figure supplement 2**. Z-AAT in a concentration-dependent manner reduces *CX3CR1* mRNA expression in PBMCs isolated from healthy donors.

**Figure supplement 3**. Z-AAT induces association of CX3CR1 with lipid rafts.

The online version of this article includes the following source data and figure supplement(s) for figure 3:

Source data 1. Source files, containing original data for Figure 3**A**,**B** to document reduced CX3CR1 surface expression monocytes (**A**) and NK cells (**B**) after treatment with Z-AAT or LPS (**A**).

**Figure supplement 1**. Gating strategy: Sequential gating to identify monocytes and NK cells from total PBMCs.

The online version of this article includes the following source data and figure supplement(s) for figure 4:

Source data 1. Source files, containing original data for Figure 4 **A**,**B** to document CD14 gene expression in PBMCs (A) and CD14 surface expression in monocytes (**B**).

**Figure supplement 1**. Inverse changes in *CD14* and *CX3CR1gene* expression in PBMCs treated with rising concentrations of Z-AAT.

## Figure supplements

**Figure 1–figure supplement 1.**
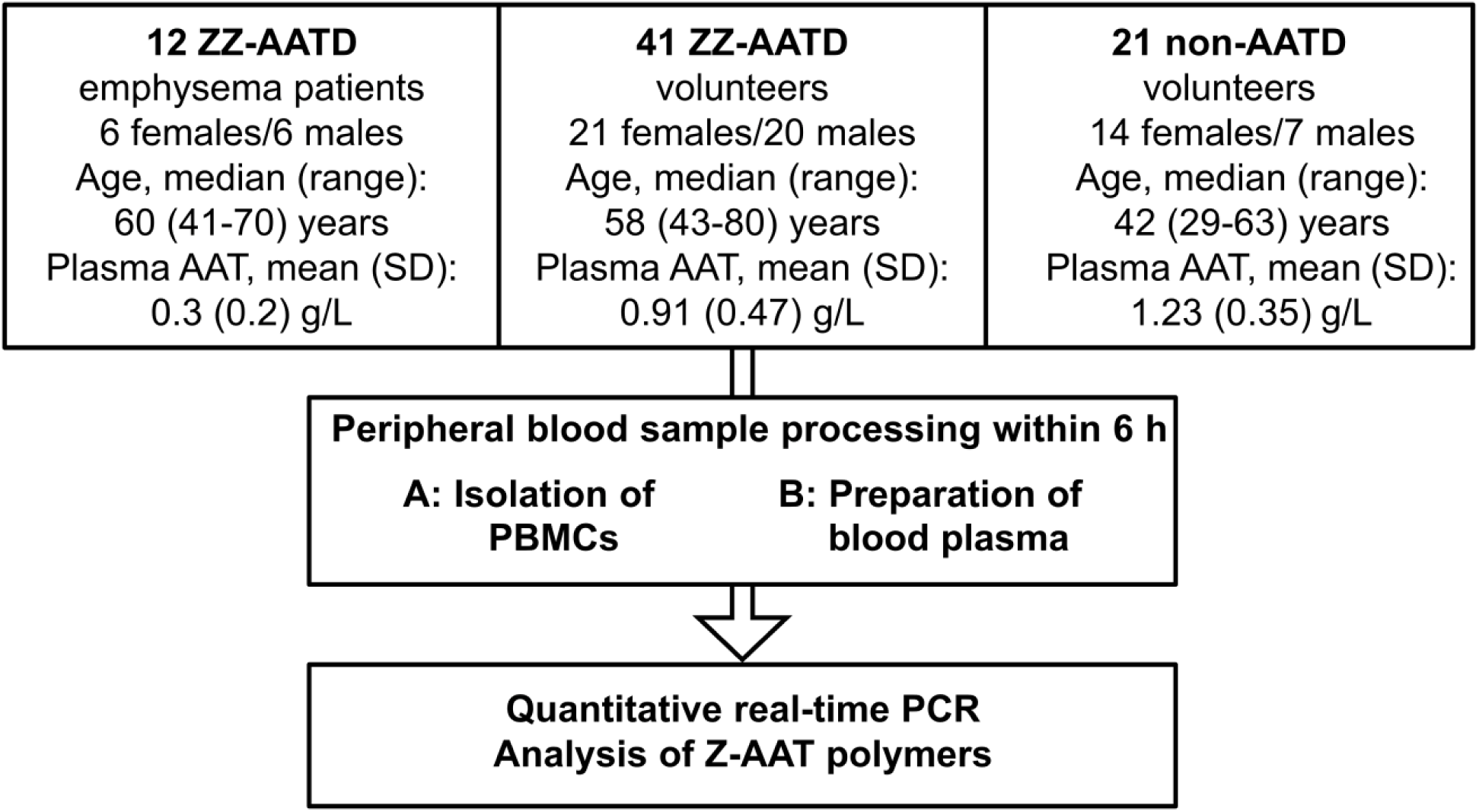
Schematic presentation of the study design. The German cohort comprises 41 ZZ-Alpha1 Antitrypsin deficient (AATD) volunteers irrespective of clinical status and medications. For control, we collected blood from 21 non-AATD volunteers with normal AAT blood levels. We prepared PBMCs for further gene expression analysis by real-time PCR and plasma for determination of Z-AAT polymers by nephelometry. The Leiden cohort consisting of 12 ZZ-AATD emphysema patients was used for gene expression analysis.

**Figure 2–figure supplement 1.**
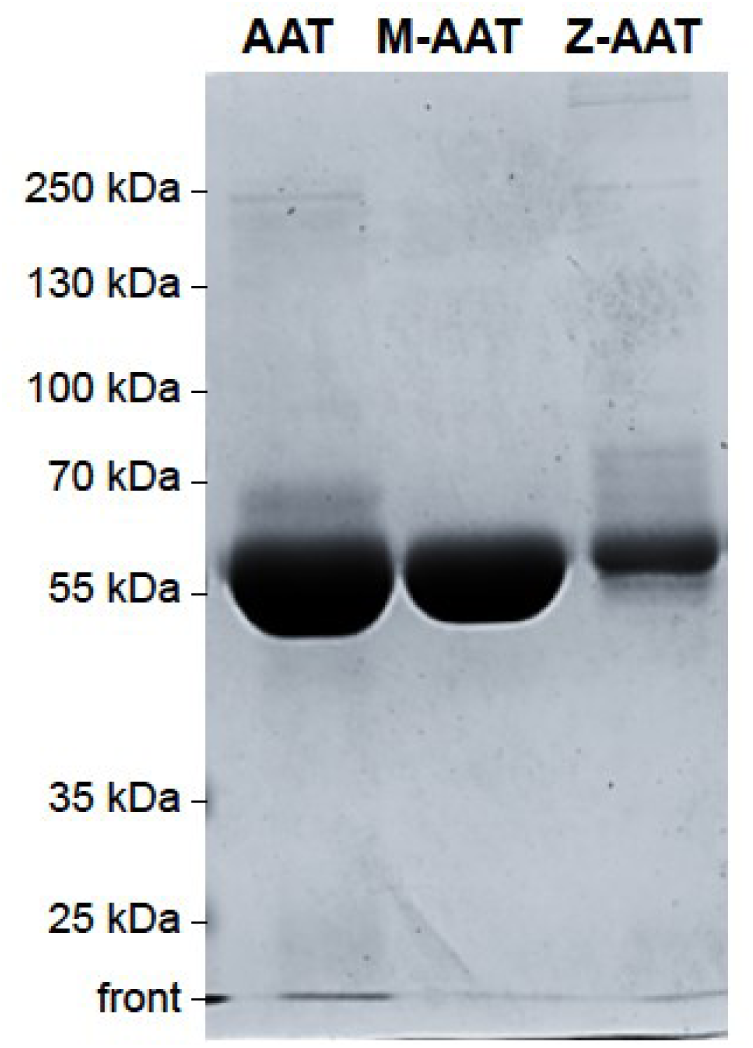
Quality control of isolated M- and Z-AAT protein by SDS-PAGE. Plasma M- and Z-AAT was purified by affinity chromatography with AAT specific Alpha1 Antitrypsin Select matrix. Proteins were run on a 10 % SDS PAA gel and stained with Coomassie Blue G250. AAT (Respreeza) was added as a positive control. Gels were run for each protein preparations. One representative gel is shown.

**Figure 2–figure supplement 2.**
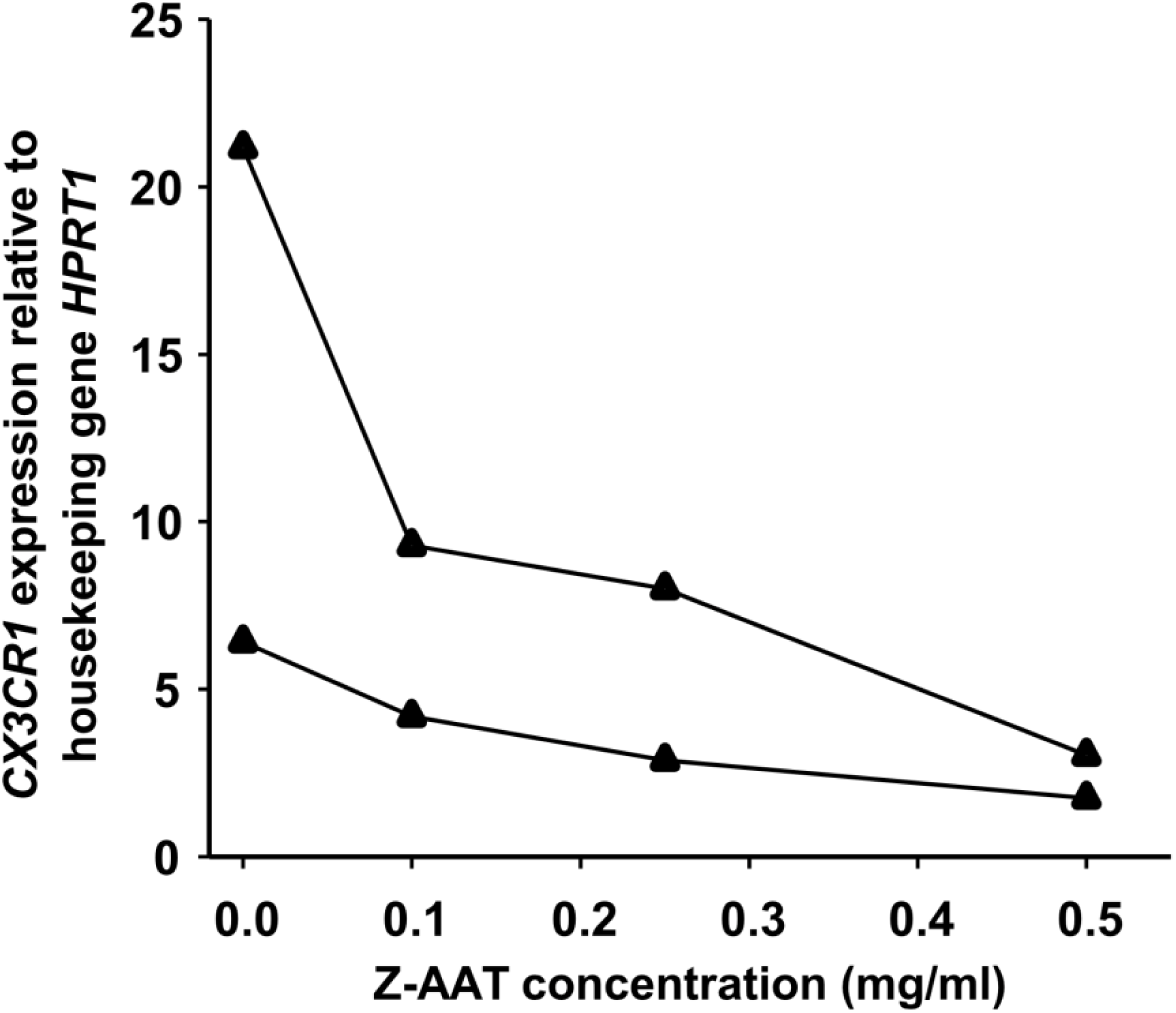
Z-AAT in a concentration-dependent manner reduces *CX3CR1* mRNA expression in PBMCs isolated from healthy donors. PBMCs were incubated for 18 h with plasma-derived Z-AAT in the concentrations as indicated. *CX3CR1* gene expression relative to *HPRT1* housekeeping gene was determined by real-time PCR using Taqman gene expression assays. Curves show results from two independent experiments. Each point represents mean of two repeats.

**Figure 2–figure supplement 3.**
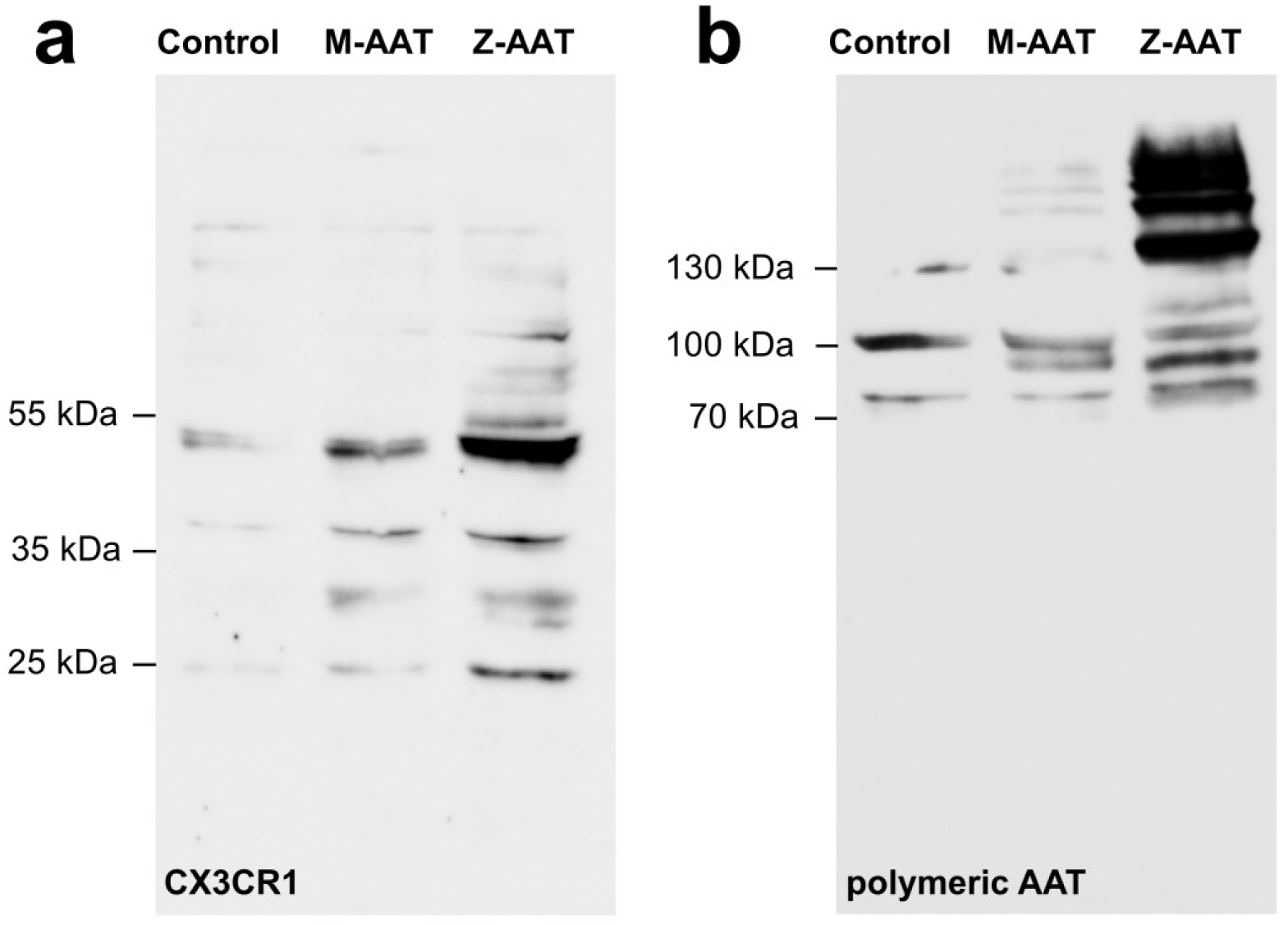
Z-AAT induces association of CX3CR1 with lipid rafts. Lipid rafts were solubilized from membrane fractions with ULTRA RIPA kit. **a** For analysis of CX3CR1, equal amounts of protein were separated by SDS-PAGE under reducing conditions. One representative blot from n = 3 independent experiments is shown. **b** For analysis of lipid raft associated AAT polymers the same samples were separated under non-reducing conditions. The Western blot was probed with monoclonal antibody (2C1) recognizing polymeric AAT. One representative blot from n = 3 independent experiments is shown.

**Figure 3–figure supplement 1.**
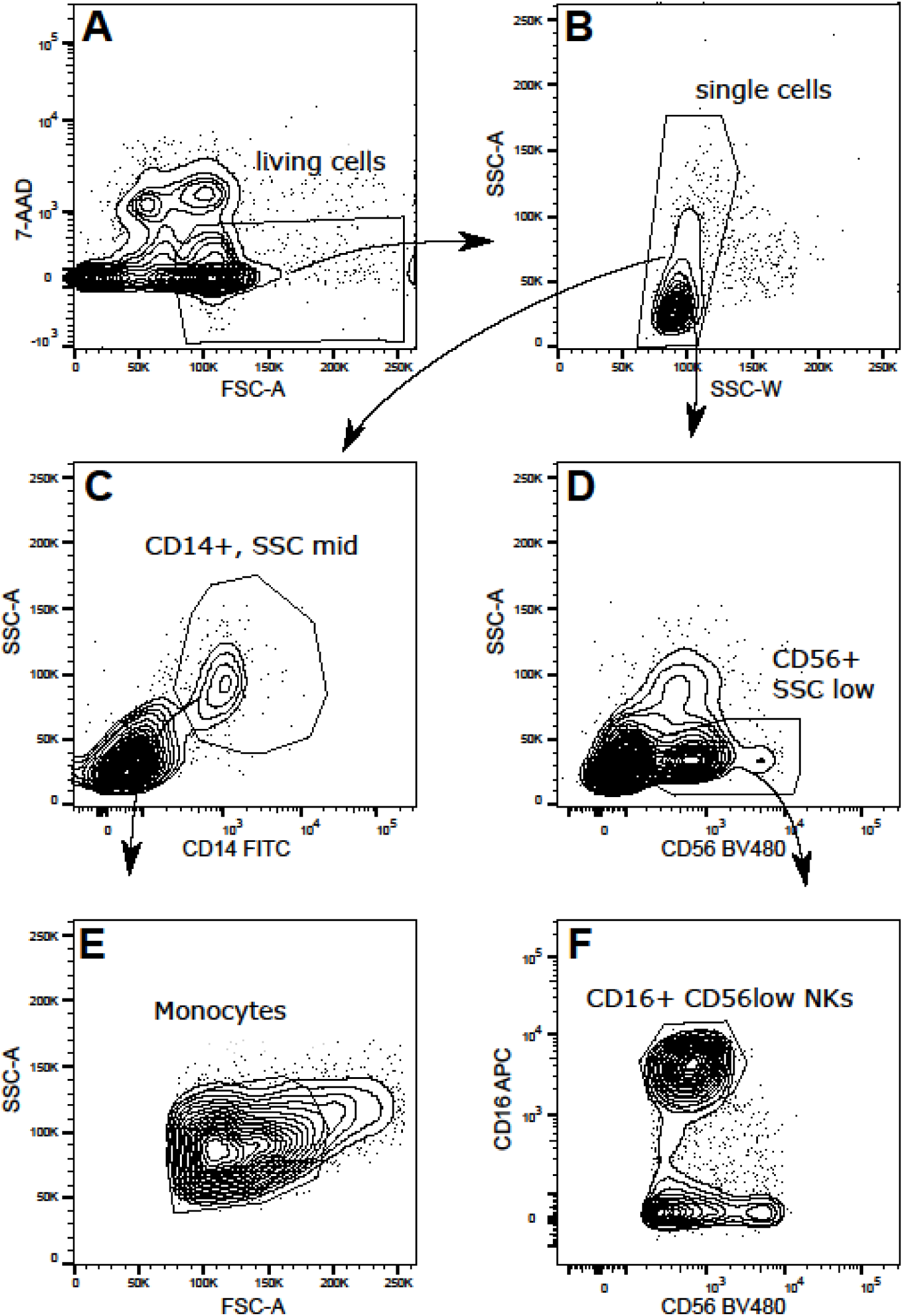
Gating strategy: Sequential gating to identify monocytes and NK cells from total PBMCs. PBMCs were stained with CD14, CD16 and CD56 in the presence of 7-amino-actinomycin D (7-AAD). Dead cells were excluded by 7-AAD. From the single cells, CD14^+^ cells with a mid-level of granularity (SSC mid) were further gated for a mid-level FSC to identify monocytes. For the identification of NK cells, CD56^+^ cells with a low-level of granularity (SSC low) were further selected by CD16. The above plots are representative for four different donors.

**Figure 4–figure supplement 1.**
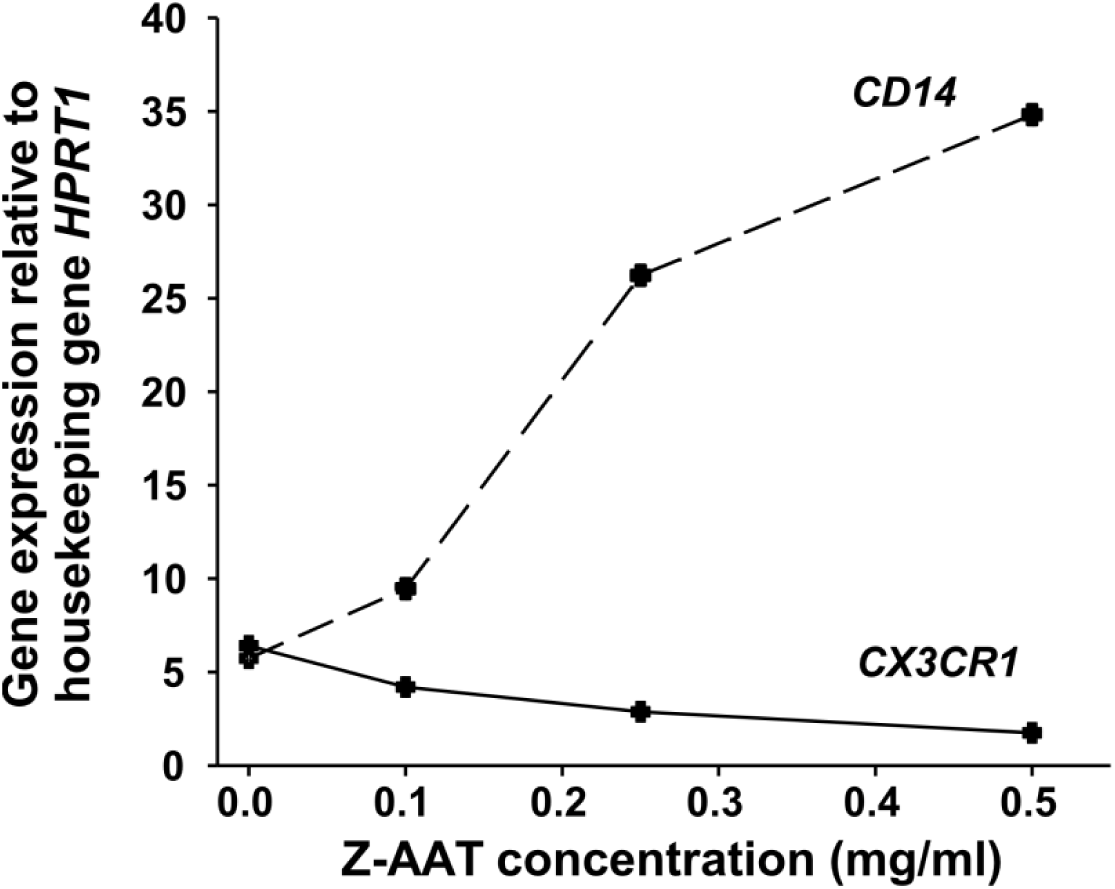
Inverse changes in *CD14* and *CX3CR1* mRNA expression in PBMCs treated with different concentrations of Z-AAT. PBMCs were incubated for 18 h with plasma-derived Z-AAT in the concentrations as indicated, or with RPMI medium alone (control). *CX3CR1* and *CD14* gene expression relative to *HPRT1* housekeeping gene was determined by real-time PCR using Taqman gene expression assays. Curves represent two independent experiments.

**Figure 4–figure supplement 2.**
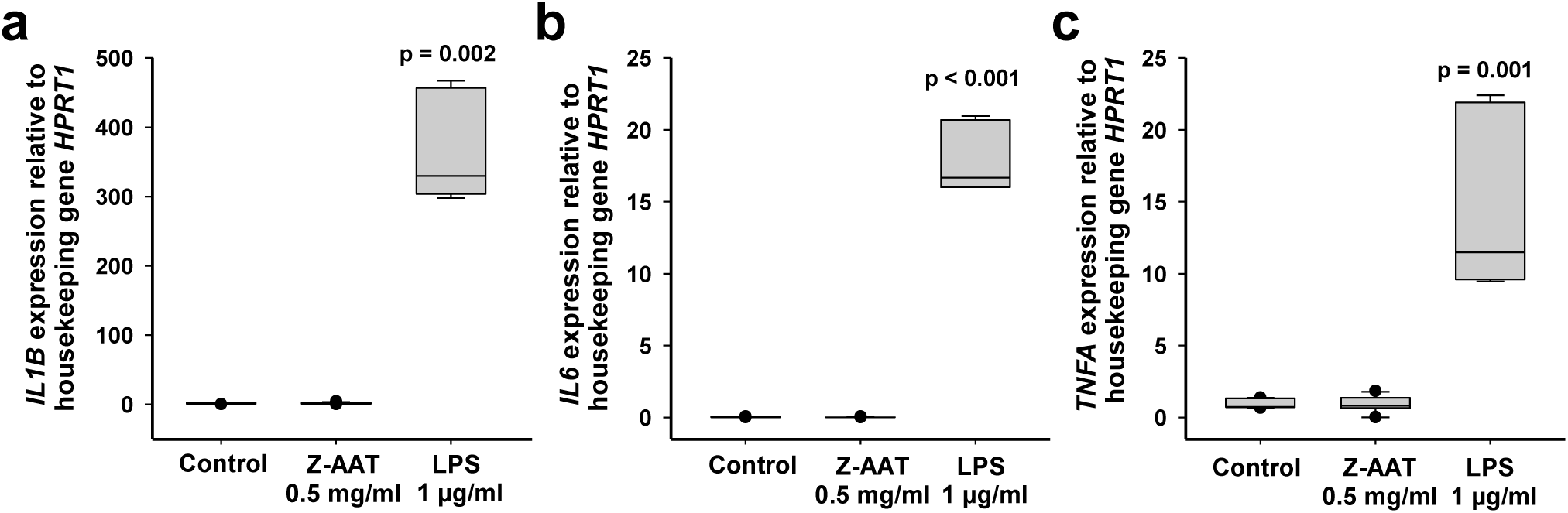
Z-AAT doesn’t induce cytokine expression. PBMCs were treated with Z-AAT or LPS (used as a positive control) for 18 h. *IL1B* (**a**), *IL6* (**b**) and *TNFA* (**c**) mRNA expression relative to *HPRT1* housekeeping gene was determined by real-time PCR using Taqman gene expression assays. Data presented as median (IQR) in box and whisker plots, lines represent medians, outliers are defined as data points located outside the whiskers. n = 4 independent experiments. p-values were calculated by Kruskal-Wallis one-way analysis.

